# Culturing of ‘Unculturable’ Subsurface Microbes: Natural Organic Carbon Source Fuels the Growth of Diverse and Distinct Bacteria from Groundwater

**DOI:** 10.1101/2020.05.01.073353

**Authors:** Xiaoqin Wu, Sarah Spencer, Sara Gushgari-Doyle, Mon Oo Yee, Jana Voriskova, Yifan Li, Eric J. Alm, Romy Chakraborty

## Abstract

The recovery and cultivation of diverse field-related microorganisms from the terrestrial subsurface environment remains a challenge despite recent advances in modern molecular technology. Here we applied natural organic carbon (C), i.e., sediment-derived natural organic matter (NOM) and bacterial cell lysate, to groundwater microbial communities for a 30-day enrichment incubation, followed by conventional direct-plating for isolation. The groundwater was collected from a background well at the Oak Ridge Reservation Field Research Center, Tennessee. As a comparison, we also included enrichments amended with simple organic C sources, including glucose, acetate, benzoate, oleic acid, cellulose, and mixed vitamins. Our results demonstrate that complex natural organic C sources are more effective in enriching diverse bacterial species from groundwater than simple organic C sources. Microcosms amended with simple organic C (glucose, acetate, benzoate, or oleic acid) show significantly lower biodiversity than unamended control and are dominated by only few phyla such as *Proteobacteria* and *Bacteroidetes*. In contrast, microcosms amended with complex natural organic C (sediment NOM or bacterial cell lysate) display significantly higher biodiversity, and enrich distinct species from the phyla that are poorly represented in published culture collections (e.g., *Verrucomicrobia, Planctomycetes*, and *Armatimonadetes*). Our subsequent isolation efforts from natural organic C-amended enrichments led to 222 purified bacterial isolates representing 5 phyla, 16 orders, and 54 distinct species including candidate novel, rarely cultivated, and undescribed organisms.

**Importance:** Innovative strategies for recovering bacterial strains representing the true diversity of microbial communities in the terrestrial subsurface would significantly advance the understanding of ecologically critical taxa residing in these ecosystems. In this study, we demonstrate that complex natural organic C that mimic the naturally available resources for microbes encourages the growth of diverse bacteria much more robustly than traditional simplistic organic C sources. Results from this study will substantially advance and improve the design of strategies to effectively cultivate and isolate diverse and novel subsurface microorganisms in the laboratory. Obtaining axenic cultures of the ‘once-unculturable’ microorganisms will greatly enhance our understanding of microbial physiology, function, and roles in different biogeochemical niches in terrestrial subsurface ecosystems.

## Introduction

Compared to animal and plant hosts, other non-human environments on Earth such as marine sediment, seawater, soil, and the terrestrial subsurface host prodigious and undiscovered microbial populations, as most of them have never been cultured and characterized in the laboratory^1^. In the terrestrial subsurface, it is estimated that there are 2,500 × 10^26^ microbial cells, of which more than 70% belong to uncultured clades and thus their physiologies and ecological impacts remain largely mysterious^1^. Despite rapid technological advances in modern molecular tools—such as metagenomics, metatranscriptomics, and metaproteomics—for identification of key microbial taxa and critical metabolic processes in a given environment, a complete interpretation of omics-based data is still constrained by the unavailability of reference genomes and isolates^2^. Challenges in microbial cultivation/isolation in the laboratory have impeded the ability of microbiologists to fully investigate the roles and function of microbes in terrestrial subsurface ecosystems.

Successful recovery and cultivation of environmental microbes in the laboratory critically depends on appropriate growth media and incubation conditions that best mimic the ecological habitat of the bacteria^3^. Enrichment culturing is a common initial step in microbial isolation to select for microorganisms with specific metabolisms within the total microbial population. The choice of organic carbon (C) substrate is of paramount importance in enrichment media composition. Yeast extract and simple organic compounds such as glucose, acetate, lactate, pyruvate, and casamino acids are amended routinely, either as an individual C source or as a mixture with the understanding that most microbes utilize these C substrates^4^. However, these labile C compounds commonly lead to selective and biased growth of microorganisms with kinetic advantages (e.g. fast growing microorganisms), generally considered as ‘weeds’^5, 6^, and have rarely recovered slow growing metabolically active microbes from the environment^7^. For this reason, despite the rapid advances in “omics” technologies, we have still only been able to cultivate less than 2% of microbes on Earth in the laboratory^8-10^.

Rationally designed growth medium that closely mimics the natural environmental habitats of microorganisms has proven to be an effective strategy in recovering diverse and previously uncultivable organisms from various environments^11-16^. Specifically, microorganisms in subsurface are reported to grow optimally in low nutrient availability or oligotrophic conditions^17^. In groundwater, natural organic matter (NOM) derived from the adjacent sediment provides the available C source for microorganisms. Our previous study shows that sediment-derived NOM contains a myriad of heterogenous organic compounds—mostly recalcitrant C such as lignin-like compounds and a small portion of relatively labile C such as carbohydrate- and protein-like compounds^18^. Other natural C source available for microorganisms in groundwater can be from dead, lysed microbial biomass turnover. Despite the potential of these natural C sources for diverse microbial cultivation under laboratory conditions, no research has been reported on the application of sediment NOM or microbial cell lysate for cultivation/isolation of microorganisms from the terrestrial subsurface environment.

In this study, we aim to develop an effective cultivation strategy using naturally occurring complex C to recover diverse, rarely cultivated, and novel bacteria from groundwater collected at the Field Research Center (FRC) in Oak Ridge, Tennessee. Our results show that natural complex C such as NOM and bacterial cell lysate are much more effective than conventional simple organic C sources in encouraging the growth of diverse and distinct bacteria from groundwater, providing a platform for the recovery of undiscovered bacteria that constitute ‘microbial dark matter’ in the subsurface. Results from this study will aid in the design of successful cultivation strategies to unlock diverse novel, previously uncultured, or ecologically important subsurface microbes for phenotypic and genomic analysis, which will greatly advance our understanding of microbial physiology, roles, and function in biogeochemical cycles in the terrestrial subsurface.

## Results

In this study, we applied a two-step workflow for cultivating and isolating a broad diversity of bacteria from groundwater. Microcosm enrichments amended with different C sources were used as the first step to enrich bacterial species from Oak Ridge FRC groundwater. We evaluated two types of complex natural organic C source: FRC sediment-extracted NOM and bacterial cell lysate. The bacterial cell lysate was prepared using a native, naturally abundant bacterial strain isolated from FRC groundwater to mimic the cell lysis products available for groundwater microorganisms. For comparison, we also evaluated several types of simple organic C source, i.e., conventional C source (glucose and acetate), naturally occurring compounds (benzoate, oleic acid, and cellulose), and mixed vitamins. The mixed vitamins were included because they are often added as supplements to bacterial growth media^19, 20^, and we wanted to test whether they are a limiting factor for support of microbial growth in this experiment. After enrichment cultivation, conventional direct plating was conducted to obtain axenic bacterial isolates from enrichment cultures amended with complex natural organic C source (i.e. NOM or cell lysate).

### Natural organic C source increases bacterial diversity in enrichments

Our results show that both C type and length of incubation have significant influence on bacterial community structure in enrichment cultures. Statistical analysis reveals that C type is the major driver of community dissimilarity (MANOVA/*adonis, R*^2^ = 0.56; ANOSIM, *R* = 0.88, *p* = 0.001), with incubation time contributing to a lesser extent to variation (MANOVA/*adonis, R*^2^ = 0.09; ANOSIM, *R* = 0.12, *p* = 0.001). We accordingly grouped samples by NMDS ordination based on the type of amended C source (Figure 1). The bacterial community in enrichments amended with glucose, acetate, benzoate, oleic acid, bacterial cell lysate, or sediment NOM clearly differs from the unamended control. The bacterial community composition in cultures amended with small organic C (glucose, acetate, benzoate, or oleic acid) are noticeably similar to each other at an early stage of incubation, and then diversify at later stages—while bacterial community composition in cultures amended with complex natural organic C (bacterial cell lysate or sediment NOM) separate far from other groups from early on.

**Figure 1.**
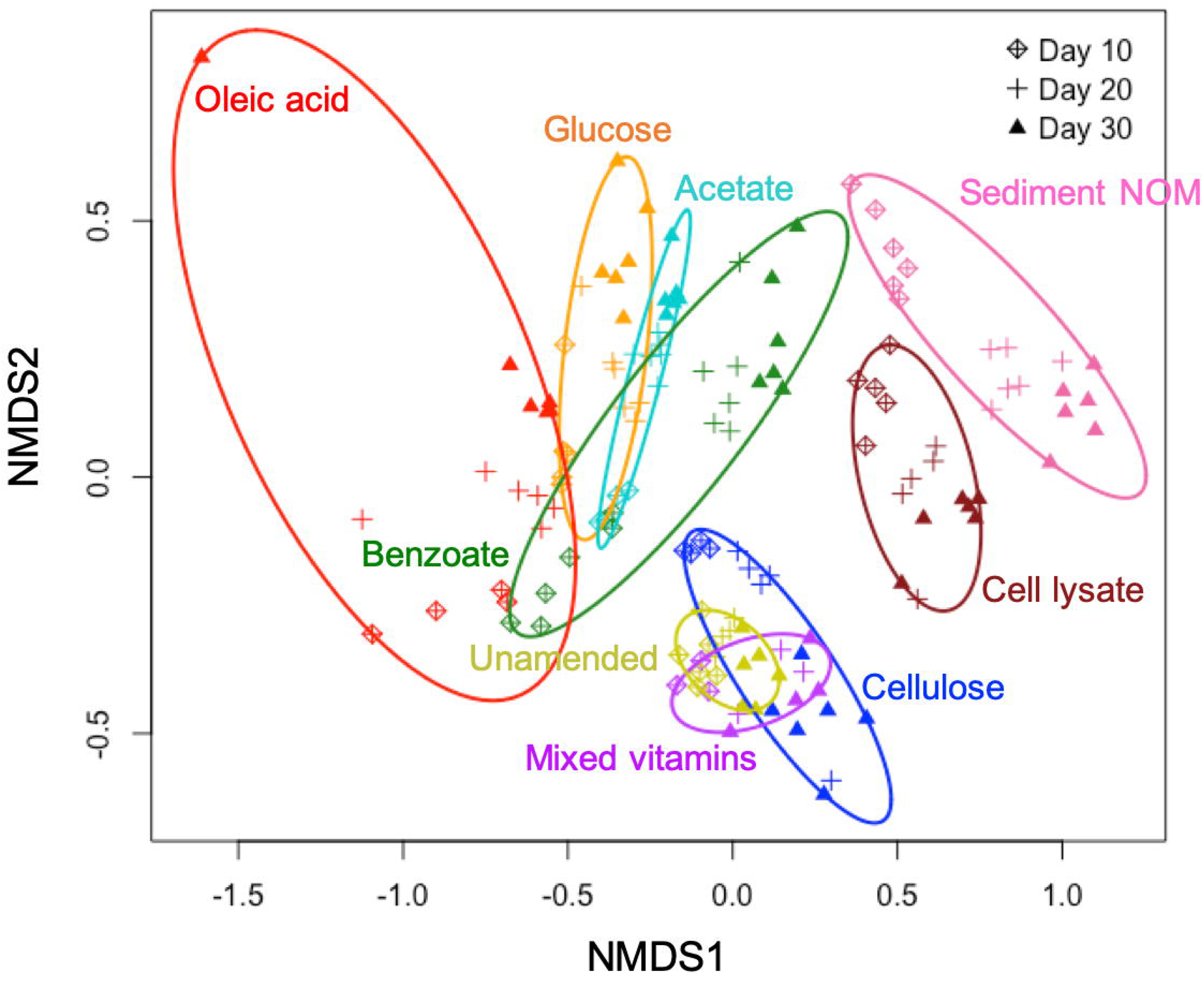
Non-metric multidimensional scaling (NMDS) based on Bray-Curtis dissimilarities of bacterial community composition.

We also observe that the diversity of the enriched bacterial community is related to the complexity of amended C substrates (Figure 2). The bacterial diversities in enrichments amended with small organic C (glucose, acetate, benzoate, or oleic acid) are generally lower than those in unamended control (Figure 2A). This strongly suggests that providing microbial communities with a simple, small organic C source in growth media will decrease diversity and lead to enrichment of a select few bacterial species that preferentially utilize these C substrates. In contrast, the bacterial diversities in enrichments amended with complex natural organic C (bacterial cell lysate or sediment NOM) are higher than those in unamended control, demonstrating the power of natural organic C in promoting growth of diverse bacteria (Figure 2A).

**Figure 2.**
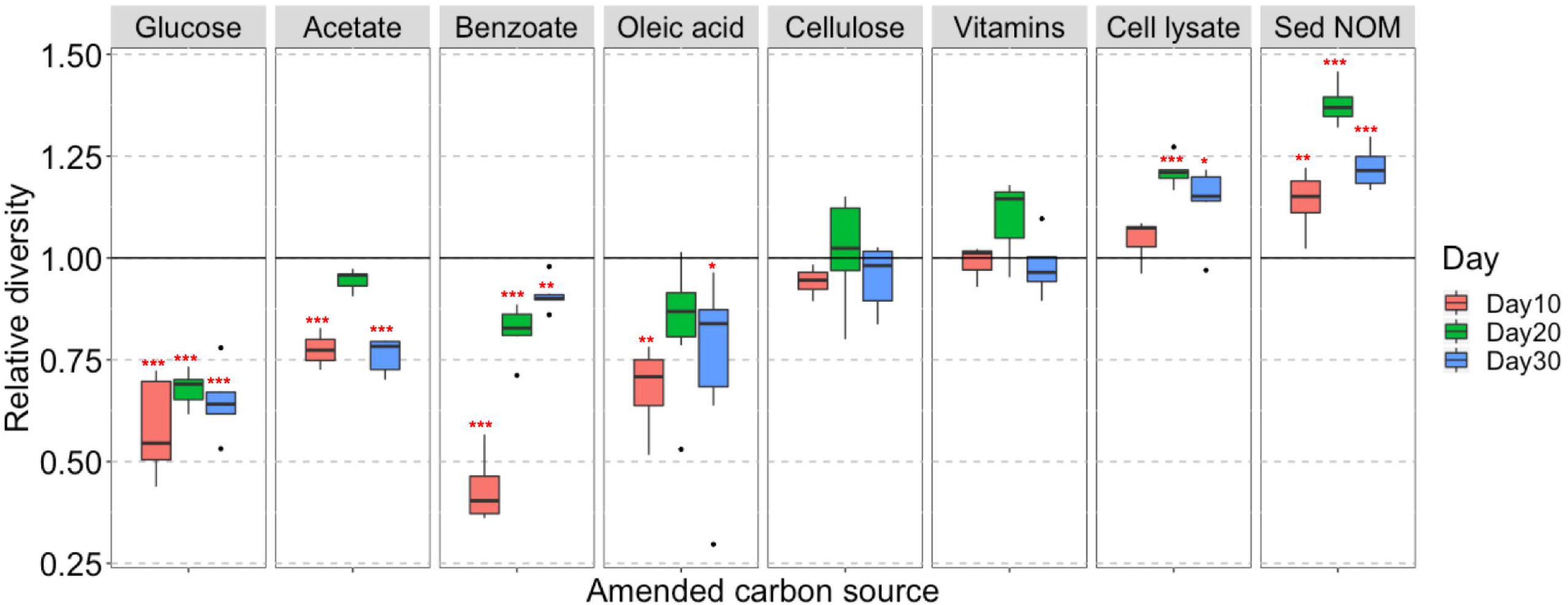
Diverse bacteria enriched with complex natural organic C sources. (A) Box and whisker plots illustrating the relative diversity of bacterial community in enrichments amended with different C sources as compared to the unamended control. Relative diversity is calculated as *H*_*i*_*’* /*H*_*0*_*’. H*_*i*_*’* is Shannon’s diversity index of individual sample, *H*_*0*_*’* is the average Shannon’s diversity index of unamended control at corresponding sampling time point. Significant differences between C-amended group and unamended control is indicated by *** when *p* < 0.001, ** when *p* < 0.01, and * when *p* < 0.05. (B) Temporal community structures of each C-amended group reported as relative abundance of taxonomic phyla over 3 timepoints (Days 10, 20, and 30).

Cellulose and mixed vitamins show relatively little influence on bacterial community composition and diversity in comparison with the unamended control (Figure 1 and 2A), and therefore are not included in our further statistical analysis.

### Natural organic C source enriches distinct bacterial taxa

We investigate the short-term response of bacterial community structure to different C sources in enrichment cultures via 16S rRNA gene survey. Out of the quality-filtered reads, organisms from 21 phyla and 94 orders are taxonomically identified, covering 71–100% of all reads, except for two samples (57% and 60%) in the bacterial cell lysate-amended group. All phyla and abundant orders (with relative abundance >1% in any sample) are presented in Figure 2B and Supplementary Figure S1, respectively. *Proteobacteria* and *Bacteroidetes* are the two most dominant phyla in all groups, especially in those amended with small organic C. It is worth noting that the phyla *Verrucomicrobia, Planctomycetes*, and *Armatimonadetes*, which are rarely cultivated from environmental samples, are enriched abundantly in sediment NOM-amended cultures with clear succession patterns (Figure 2B). The relative abundance of *Verrucomicrobia* significantly diminishes over time from 18–27% at Day 10 to less than 2% at Day 30. Meanwhile *Planctomycetes* becomes one of the major phyla at later stages, with relative abundance increasing from 0.1–1% at Day 10 to 5–33% at Day 30. *Armatimonadetes* also increases during the incubation period, with relative abundance up to 10% at Day 30.

In microcosms amended with small organic C, only a few taxonomic orders such as *Caulobacterales, Burkholderiales, Rhodocyclales*, and *Cytophagales* are enriched, while in complex natural organic C-amended microcosms, diverse taxonomic orders are enriched, including those scarcely enriched in other groups, e.g., *Sphingobacteriales, Gemmatales, Planctomycetales, Verrucomicrobiales*, and *Solibacterales* (Supplementary Figure S1).

Based on one-way ANOVA with Dunnett’s test results, we identify a total of 166 OTUs that are promoted to grow by test C sources, with significantly (*p* < 0.01) increased relative abundances in C-amended enrichment cultures compared to the corresponding unamended control at each time point. These promoted OTUs widely distribute across 11 phyla (Figure 3A). Compared to simple organic C sources (glucose, acetate, benzoate, and oleic acid), the complex natural organic C sources (sediment NOM and bacterial cell lysate) show a great advantage in promoting the growth of diverse and distinct bacterial species. Most of promoted OTUs (110 out of 166) are exclusively promoted by complex natural organic C sources (Figure 3B), especially those from rarely cultured phyla *Verrucomicrobia, Planctomycetes*, and *Armatimonadetes*. A small portion (41 out of 166) are exclusively promoted by simple organic C sources, most of which are from the phyla *Proteobacteria* and *Bacteroidetes*, and a few from *Acidobacteria* and WPS-2 (Figure 3B). There are 15 OTUs that can be promoted by both simple and complex organic C sources, suggesting that they likely harbor the metabolic potential for utilizing diverse C sources, from simple organic C to complex natural organic C.

**Figure 3.**
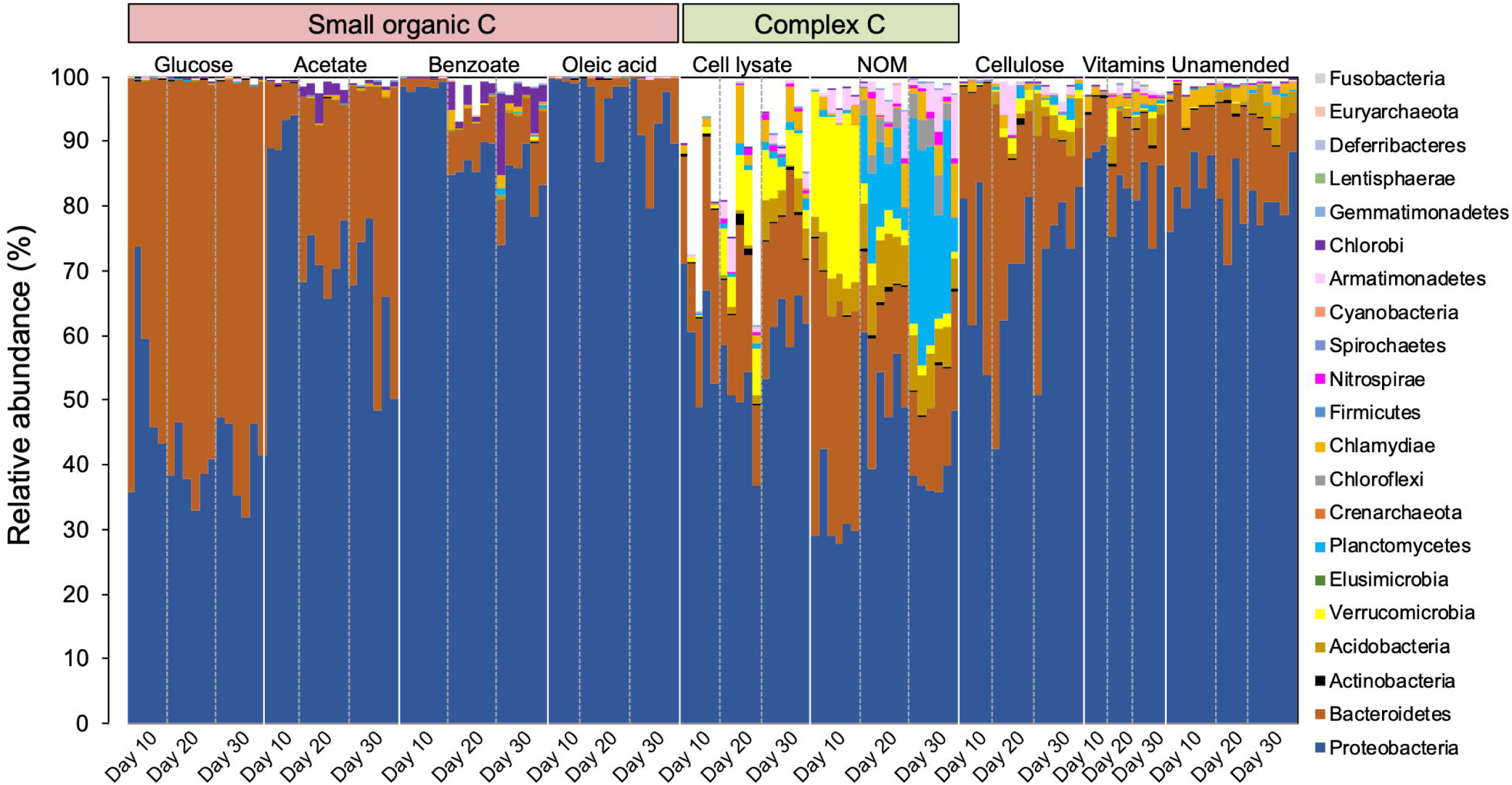
Selected bacterial species (one-way ANOVA with Dunnett’s multiple comparison adjustment, *p*-value < 0.01) that are promoted to grow by simple C sources (glucose, acetate, benzoate, and oleic acid) and complex natural organic C sources (bacterial cell lysate and sediment NOM) in enrichment cultures as compared to the unamended control. (A) Phylogenetic tree (constructed with RAxML) of ‘promoted’ OTUs. OTU labels are highlighted according to their phylum. Bold OTU labels designate OTUs for which we isolated representative species (99-100% identity). Markers surrounding the tree denote the day (10 - square, 20 - circle, or 30 - star) and C substrate in which the OTU is significantly enriched. (B) The number of OTUs that are promoted exclusively by simple C sources, exclusively by complex natural organic C sources, and by both simple and complex C sources.

Besides amended C source, incubation time also affects the enriched bacterial species. In those 110 OTUs exclusively promoted by complex C sources, we observe slow growers (25 out of 110) that exhibit significant enrichment at late incubation stage (on Day 30), and also consistent growers (29 out of 110) which are enriched in the cultures consistently from Day 10 to Day 30 (Figure 3A).

### Novel bacterial isolates from natural organic C-amended enrichments

Since complex natural organic C shows greater potential in enriching diverse and distinct bacterial species, we then use the complex C (bacterial cell lysate and sediment NOM)-amended enrichments as inocula in our further isolation work. In this study, we obtained a total of 222 bacterial isolates (Supplementary Table S2) representing 5 phyla, 16 orders, and 54 distinct species (Figure 4 and Supplementary Table S3). A comparison between the enrichment and isolation results shows that our bacterial isolates represent one-third (10 out of 33) of the enriched orders (Supplementary Figure S1), and 16% (27 out of 166) of the promoted OTUs (Figure 3A and Supplementary Table S1). We obtained representative isolates not only for the OTUs that are exclusively promoted by complex C source (e.g., denovo1156), but also for the OTUs that are exclusively promoted by simple C sources and exist at very low relative abundance (< 0.1%) in the complex C-amended enrichment cultures (e.g., denovo2244). We also obtained bacterial isolates representing slow growers (e.g., denovo3150) and consistent growers (e.g., denovo243, denovo422, denovo1156, and denovo2687) from our isolation efforts (Figure 3A).

**Figure 4.**
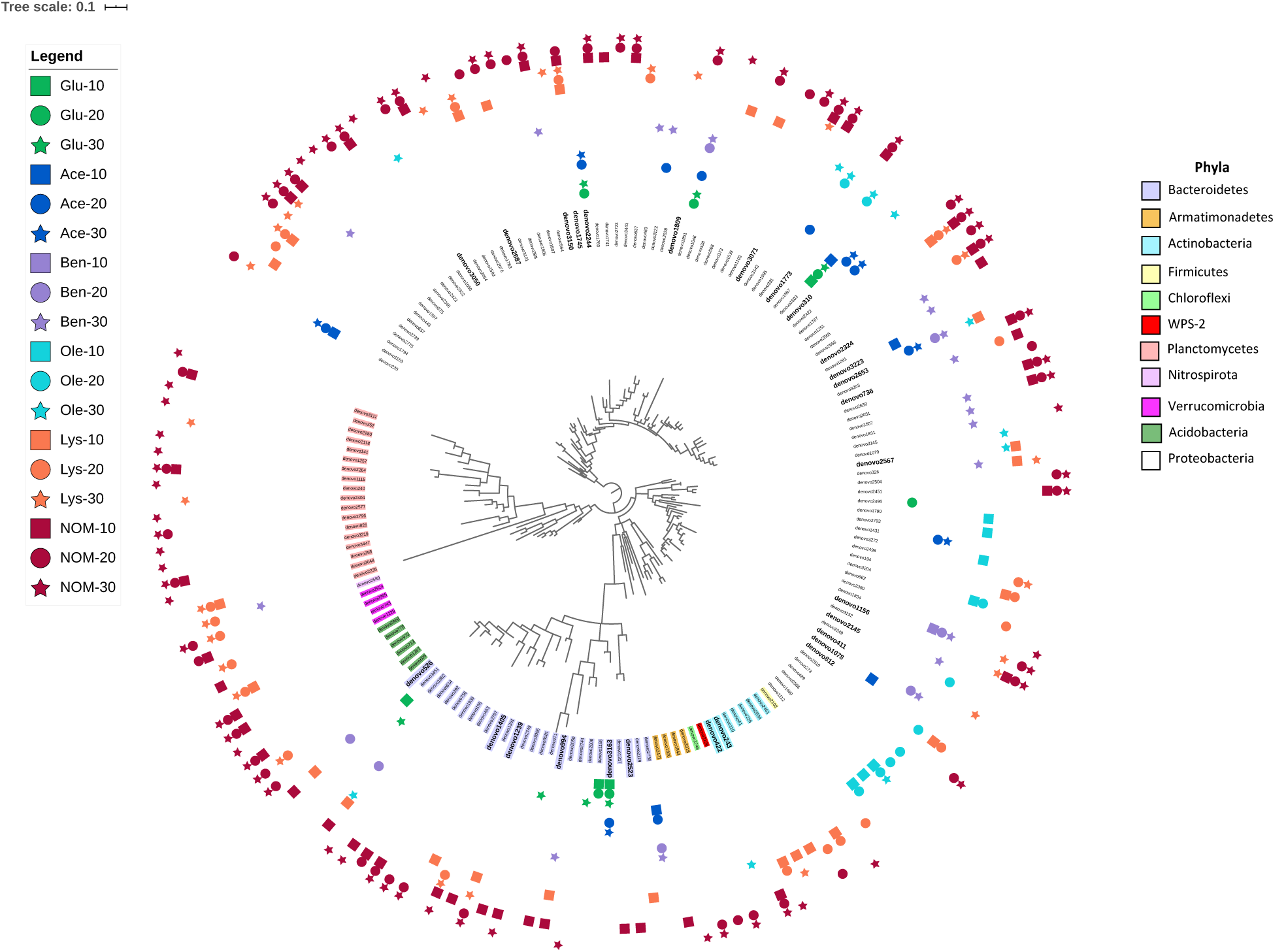
Phylogenetic tree of representative bacterial isolates representing 54 distinct species. The tree is constructed from near full-length 16S rRNA gene sequences. Undescribed species (black), novel candidate species (blue), and novel candidate genus members (red) are shown by dot colors. The class and order of the representative isolates are listed on the right.

Thresholds for determining the novelty of an isolate based on 16S rRNA gene sequence similarity have differed slightly in different reports^21, 22^. Here we apply the thresholds of 98% for novel species, 95% for novel genera, and 90% for novel families^15^. According to these criteria, of the 54 distinct bacterial species isolated from FRC groundwater, nine belong to candidate novel species and three belong to candidate novel genera (Figure 4 and Supplementary Table S3). These novel isolates distribute across two phyla *Proteobacteria* and *Bacteroidetes*, which are the most dominant phyla in the original FRC groundwater sample (data not shown) as well as enrichment cultures in this study. Besides, there are 9 undescribed species unassigned at the genus level in the SILVA database, indicating that they are from the less characterized genera with unresolved taxonomy. The reconstructed phylogenetic trees for these novel and undescribed organisms are presented in Figure 5 and Supplementary Figure S2.

**Figure 5.**
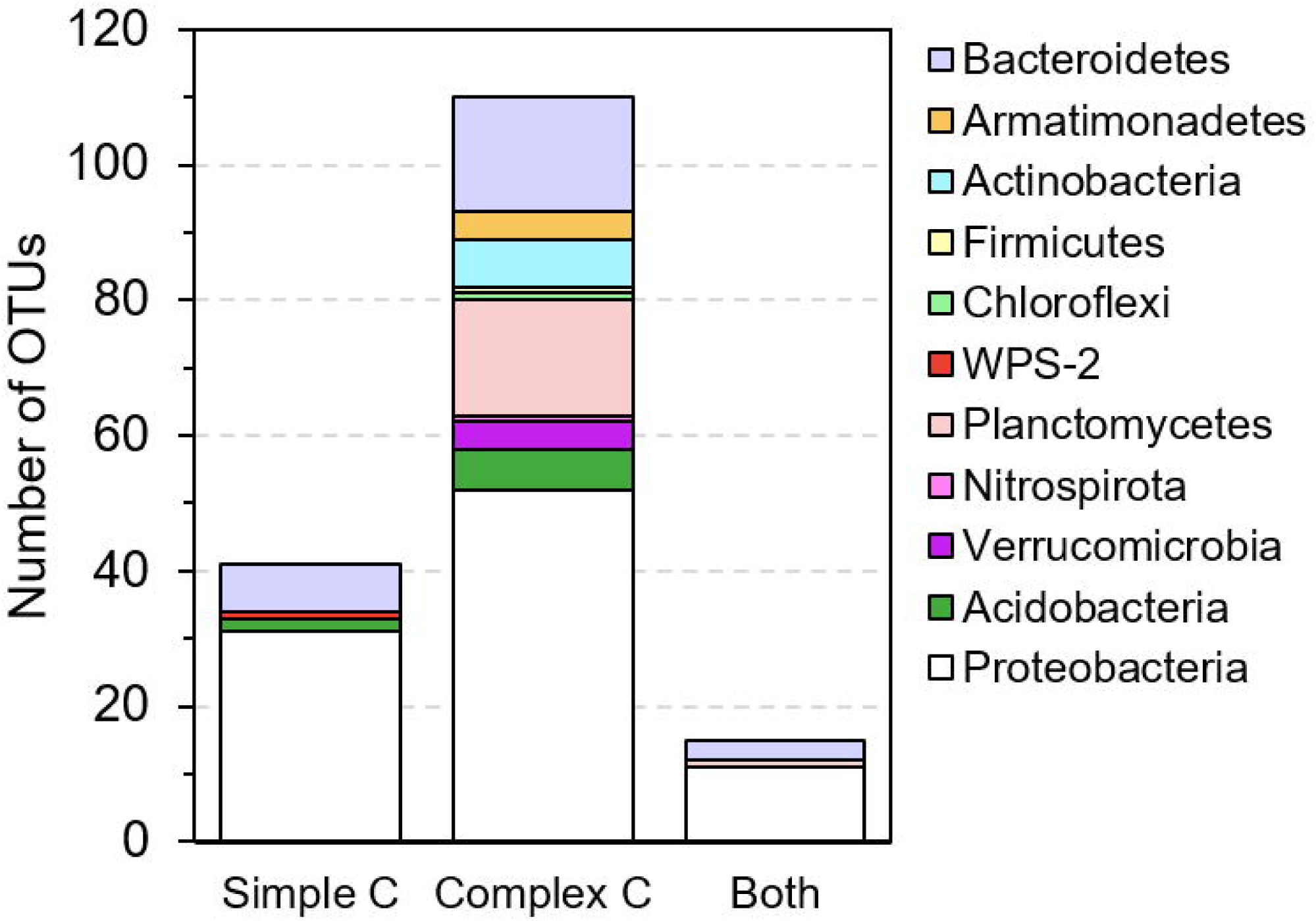
Phylogenetic tree of FW305-C-21 and the most similar bacteria based on 16S rRNA genes. Scale bar indicates a change of 0.02 per nucleotide. The 16S rRNA gene sequences were aligned using SINA against the SILVA alignment and the maximum likelihood tree was calculated using RAxML. The top five organisms are as far cultured organisms within this family.

## Discussion

There is a compelling need for improving the recovery of diverse bacteria from environments. Several ongoing efforts include modification of growth media/conditions^23^, use of diluted medium or serial dilution culture^24, 25^, and cultivation with physical separation (e.g., iChip^26^ or diffusion chambers^12, 27, 28^). However, the collective capability for recovering microorganisms from the terrestrial subsurface, especially those mediating critical biogeochemical cycles, is still limited. This bottleneck continues to hinder a thorough investigation of microbial ecology and understanding of physiology and true metabolic potential of key organisms residing in subsurface ecosystems.

In this study, we demonstrate that natural organic C sources (sediment NOM and bacterial cell lysate) fuel the growth of much more diverse and distinct groups of microbes compared to traditional simple organic C sources. Natural organic C is a mixture of heterogeneous naturally occurring substrates^18^, making it a suitably appropriate C source in encouraging growth of diverse microbe representatives of in-situ environmental communities and typically those not cultivated in the laboratory. As shown in our enrichment results, almost all species from the rarely cultured phyla *Verrucomicrobia, Planctomycetes*, and *Armatimonadetes* show exclusive preference for complex C source especially sediment NOM (Figure 3A, 3B). To date, only a handful of *Verrucomicrobia* isolates have been successfully cultivated^24, 29-33^, although members of this bacterial phylum are highly prevalent in the environment^34, 35^. It is reported that only ∼2% of strains in *Planctomycetes* have been isolated in pure cultures^36^. *Planctomycetes* are of deep interest to microbiologists because of their unique characteristics. They are reported to be comparatively slow growing organisms with low demand for C and nitrogen sources^36^, which may explain their significant enrichments at late incubation stage (Day 20 and 30) in this experiment (Figure 2B). The phylum *Armatimonadetes* lacked an isolated representative until 2011^37^, and so far, only a few cultivated strains in this phylum have been reported^37-40^. In this study, we observe that most of promoted *Planctomycetes* and *Armatimonadetes* species are exclusively enriched in sediment NOM-amended cultures only at late incubation stage (Figure 3A), indicating that these slow growers may possess high metabolic potential of utilizing relative recalcitrant C in sediment NOM, therefore avoid competition for labile C in sediment NOM with competitive fast growers^18^. Although we have not yet isolated pure bacterial strains from *Verrucomicrobia, Planctomycetes*, and *Armatimonadetes*, the results of this study will augment our ongoing efforts to obtain pure isolates in the very near future.

Applying the two-step cultivation strategy, i.e., enrichment followed by isolation, we obtained pure cultures of 54 distinct bacterial species from groundwater, some of which are novel, previously uncultured, and uncharacterized organisms. Notably, we obtained two similar isolates (FW305-C-21 and FW305-C-23) from the candidate family env.OPS 17, which is a poorly described family in literature and lacks representative isolates. To date, there are only five described cultured organisms within this family, found to be associated with ascomycetous ectomycorrhizal fungi^41^ or in freshwater springs (NCBI database). Our isolates FW305-C-21 and FW305-C-23 are distinct from those five cultured organisms (Figure 5, only showing FW305-C-21). While several of their close neighbors have been detected via molecular tools in various environments including pit^42^, drinking water^43^, uranium mining wastes^44^, freshwater lake, pond, soil, and sludge (information from the NCBI database), our isolates are the very first cultured organisms in this distinct clade. Our enrichment results show that species from this candidate family env.OPS 17, i.e., denovo1405 (with representative isolate FW305-C-21 and FW305-C-23) and denovo2797, exclusively prefer complex natural C sources (Figure 3A and Table S1), which may explain why these organisms have rarely been cultivated in the laboratory.

We also obtained pure cultures of three distinct species: FW305-C-2, FW305-C-3, and FW305-C-57, from an undercharacterized order *Salinisphaerales* which has only 17 reported genomes so far, the second-fewest in the class *Gammaproteobacteria* (NCBI lifemap). The isolates FW305-C-2 and FW305-C-3 are novel candidate genus members (Figure 4 and Table S3). Phylogenetic analysis of the isolate FW305-C-3 shows that it is close to the genus *Fontimonas* (Figure S2). The isolate FW305-C-2 clusters together in the phylogenetic tree with multiple uncultured organisms (Figure S2). The only cultured organism in this clade, *Sinobacteraceae* bacterium MG649968.1, was reported very recently from surface freshwater^45^. We have therefore made a good contribution to the number of representative cultured organisms in this distinct clade.

This study demonstrates the potential of complex natural organic C, especially NOM, for enriching diverse and ecologically relevant bacterial taxa, and for retrieving pure cultures of novel, previously uncultured organisms from the terrestrial subsurface. Our cultivation strategy will benefit future development of effective and ecologically relevant cultivation/isolation strategies. These improved capabilities will be crucial for further understanding of bacterial physiology, functions, and roles in biogeochemical cycles in terrestrial subsurface ecosystems.

## Materials and Methods

### Preparation of C stock solutions

Glucose, sodium acetate, sodium benzoate, cellulose, oleic acid, vitamins, and thioctic acid were purchased from Sigma-Aldrich (St. Louis, MO). Stock solutions of glucose, sodium acetate, and sodium benzoate were prepared by dissolving the chemical in MilliQ-water (18.2 MΩ·cm, 0.22 μm membrane filtered) at 200 mM, 200 mM, and 50 mM, respectively, followed by filter-sterilization with a filtration system (0.22 μm pore-sized, polyethersulfone (PES), Corning). Oleic acid and cellulose were added to MilliQ-water at an initial concentration of 50 g/L and 20 g/L, respectively, followed by sterilization using an autoclave. Since oleic acid and cellulose are generally insoluble, their concentrations in water are expressed as initial grams per liter. A stock solution of mixed vitamins, including vitamin B_1_, B_2_, B_3_, B_5_, B_6_, B_7_, B_9_, B_10_, B_12_, and thioctic acid, was prepared in MilliQ-water according to the recipe reported by Balch et al.^19^ (Supplementary Table S4), and then filter-sterilized (0.22 μm pore-sized, PES, Corning).

Preparation of cell lysate solution was modified based on published methods^46, 47^. A strain of *Pseudomonas* spp previously isolated from Oak Ridge FRC groundwater was used for this purpose. The isolate, which by 16S rRNA gene sequence analysis was 99% identical to *Pseudomonas fluorescens*, was grown in a Luria broth (LB) liquid medium at 30°C aerobically until early stationary phase. A 30 ml aliquot of the culture was harvested, followed by centrifugation at 6,000 *g* for 20 min. The supernatant was removed, and the pellet was washed by MilliQ-water three times before being re-suspended in 10 ml of MilliQ-water. A two-step lysis procedure was used, including autoclaving and sonication in a water bath for two hours, followed by centrifugation at 6,000 *g* for 20 min. The supernatant was decanted and filtered through a syringe filter (0.2 μm pore-sized, PES, Thermo Scientific). The filtrate was stored at 4°C until use. Total organic C (TOC) content of the filtrate, i.e., cell lysate stock solution, was 2.67 g/L, measured by TOC-5050A Total Organic Carbon Analyzer (Shimadzu, Japan).

The sediment used for NOM extraction was collected from a background well FW305 at ORR-FRC, at a depth of 1.1 m below ground surface. The water-soluble fraction of sediment NOM was extracted according to a method previously developed in our lab^18^. Briefly, the freeze-dried sediment sample was extracted with Milli-Q water via rotary shaking (170 rpm) overnight at 35°C, and then sonicated in a water bath for 2 hours. The ratio of water and sediment was 4:1 (w/w). After extraction, the water-sediment slurry was centrifuged at 6000 *g* for 20 min. The supernatant was decanted and sterilized using a filtration system (0.22 μm pore-sized, PES, Corning). Filtrate containing water-extractable NOM was freeze-dried, and the lyophilized material was stored at −20°C until use.

### Microcosm enrichment

The groundwater sample was collected from a background well, which adjoined the sediment well FW305 at ORR-FRC. The sample was shipped immediately to the lab after collection with ice packs and stored at 4°C for up to 1 week. At the time of sampling, groundwater temperature was measured to be 15.4°C, pH was 6.37, dissolved oxygen (DO) was 1.39 mg/L, TOC was 5.9 mg/L, NO_3_^-^ was 0.34 mg/L, PO_4_^3-^ was less than 3.0 mg/L. The DO in groundwater exceeded 0.5 mg/L, indicating that the groundwater sample’s redox state was oxic (Ohio EPA, http://epa.ohio.gov/Portals/28/documents/gwqcp/redox_ts.pdf).

Microcosm incubation experiments were performed in pre-sterilized 250 ml-flasks, each containing 89 ml of filtered groundwater (0.22 µm pore-sized, PES, Corning) as culture medium, 10 ml of unfiltered groundwater (cell density: 2.1 × 10^6^ cells/ml) as inoculum, and 1 ml of individual C stock solution. For oleic acid and cellulose, the stocks were shaken thoroughly to mix and homogenize the solution before adding to the enrichments. For the sediment NOM-amended group, the lyophilized NOM material was fully dissolved in filtered groundwater at 200 mg/L, and filter-sterilized (0.22 µm pore-sized, PES, Corning). TOC content of the filtrate was measured to be 48.4 mg/L. A 90 ml aliquot of the filtrate (containing sediment NOM) was added with 10 ml of unfiltered groundwater. The organic C content added to the microcosms were designed to be at least five times higher than that in background groundwater (TOC 5.9 mg/L). The final concentration of substrates in the microcosms are listed in Supplementary Table S5.

An unamended control without any additional C source was included in this study, for which each flask only contained 90 ml of filtered groundwater and 10 ml of unfiltered groundwater. All groups were performed in six replicates. One blank control (without inoculum) was included in each group to monitor potential microbial contamination during incubation. All microcosms were incubated aerobically at 25°C in the dark for up to 30 days, with rotary shaking at 100 rpm. At each sampling time point (days 10, 20, and 30), a 10 ml aliquot of subculture was sampled using a sterile volumetric pipette. Microbes were concentrated by filtration through a membrane filter (0.2 µm pore-sized, PES, 25 mm, Sterlitech Corp.). The filter was then removed from the syringe filter holder and kept frozen at −80°C until DNA extraction.

### DNA extraction for microbial community analysis

Before DNA extraction was conducted, the filters were cut into 2 mm-wide stripes using sterile blades and put into DNA extraction tubes provided in PowerMax Soil DNA Isolation Kit (MO BIO Laboratories, Inc., Carlsbad, CA). DNA was extracted following the manufacturer’s protocol, and quantified using the Qubit dsDNA HS Assay Kit (Life Technologies, Eugene, OR) with a Qubit fluorometer (Invitrogen, Eugene, OR). Extracted DNA were stored at −20°C until further processing.

### 16S rRNA gene amplicon library preparation

For analysis of bacterial community composition, a two-step PCR protocol was performed. In the first step the 16S rRNA gene of V4 variable region was amplified and in the second step Illumina barcodes and adapters for sequencing were added. Extracted DNA from enrichments were each aliquoted into one of three randomized plate layouts in a laminar flow hood.

Before the first step PCR, all samples were subjected to a qPCR at multiple dilutions to determine target dilutions and threshold cycles for the first step. We used 16S rRNA gene primers PE16S_V4_U515_F and PE16S_V4_E786R (Supplementary Table S6). Both 1:1 and 1:10 dilutions of each sample were prepared in duplicate with 0.5X SYBR Green I nucleic acid gel stain (Sigma-Aldrich, St. Louis, MO), plus 280 nM for each primer and the standard reagents in the Phusion High-Fidelity PCR Kit (New England BioLabs, Ipswich, MA). Samples were then cycled under the following qPCR conditions: 98°C 30 sec; 30 cycles of 98°C 30 sec, 52°C 30 sec, 72°C 30 sec; 4°C hold. Threshold cycles were calculated and dilutions were prepared to normalize samples and ensure consistent amplification cycles across plates. PCR under the same conditions, minus the SYBR Green, was completed in quadruplicate for each sample, then quadruplicate sets were pooled and purified with Agencourt AMPure XP Beads according to the manufacturer’s protocol (Beckman Coulter, Brea, CA).

The second step PCR was used to add sample indices and final Illumina adaptors to the 16S rRNA gene amplicons. Reactions were compiled using the Phusion High-Fidelity PCR Kit according to the manufacturer’s instructions, with 420 nM indexing primers PE-III-PCR-F and PE-IV-PCR-R (Supplementary Table S6), then cycled under the following conditions: 98°C 30 sec; 7 cycles of 98°C 30 sec, 83°C 30 sec, 72°C 30 sec; 4°C hold. Final libraries were purified with Agencourt AMPure XP Beads according to the manufacturer’s protocol, then quantified and pooled prior to 2 × 250 paired-end sequencing on an Illumina MiSeq. Data are available on the NCBI database under the accession code PRJNA524696.

### 16S rRNA gene amplicon data processing and operational taxonomic unit (OTU) analysis

Raw reads were quality filtered and clustered into operational taxonomic units (OTUs) primarily with the QIIME software package^48^ using default parameters unless otherwise noted. Paired-end reads were joined with the join_paired_ends.py command, then barcodes were extracted from the successfully joined reads with the extract_barcodes.py command (and additional parameters -c barcode_in_label, -l 16, -s ‘#’). Quality filtering was accomplished with split_libraries_fastq.py (--barcode_type 16, --min_per_read_length_fraction 0.40, -q 20, --max_barcode_errors 0, -- max_bad_run_length 0, --phred_offset 33). We checked for the correct forward and reverse primers with a custom script and exported reads with primers removed and length trimmed to 225 bp. Finally, chimeric sequences were removed using identify_chimeric_seqs.py (-m usearch61, --suppress_usearch61_ref), followed by filter_fasta.py.

After quality filtering, reads were clustered into 97% OTUs, classified against a 16S rRNA database, and aligned in order to build phylogenetic trees. We ran the QIIME commands pick_otus.py, pick_rep_set.py (-m most_abundant), and make_otu_table.py to produce the OTU table. The RDP classifier was used to assign taxonomy with default parameters and the 16S rRNA training set^49^.

In this study, 16S rRNA gene amplicon sequencing resulted in over 10 million prokaryotic 16S rRNA gene reads, which were clustered into 3463 OTUs. Only rarefied OTU richness was considered further, in order to compensate for differences in sequencing depth between 144 samples. No DNA was detected in blank controls, suggesting that microbial contamination was negligible during incubation.

### Bacterial isolation

The natural complex C (bacterial cell lysate and sediment NOM)-amended enrichments at each time point were used as inocula for further bacterial isolation. Isolation from simple C enrichments was not pursued. The enrichment inoculum was streaked on a complex C agar plate, prepared using the same medium as corresponding liquid enrichment (cell lysate or NOM) with 1.5% agar (BD Biosciences, USA) and were incubated at 27°C in the dark. We also streaked the enrichment sample on diluted, commercially available culture media (with 1.5% agar), i.e., 1/25 LB, 1/25 tryptic soy broth (TSB), and 1/10 Reasoner’s 2A (R2A), to obtain as many colonies as possible. Bacterial colonies were streaked again if necessary until single colonies were obtained. The single colony was picked from a plate and transferred to 3 ml of corresponding liquid media. The liquid cultures were incubated at 27°C in the dark for up to 3 weeks before DNA extraction. All the bacterial isolates were confirmed for growth and maintenance on easily available commercial media (1/25 LB, 1/25 TSB or 1/10 R2A).

### Species identification

Genomic DNA of bacterial isolates were extracted using a PureLink Genomic DNA Mini Kit (Invitrogen, United States), following the manufacturer’s protocol. 16S rRNA genes were amplified (initial denaturation step at 98°C for 5 min, followed by 30 cycles at 95°C for 30 s, 50°C for 30 s and 72°C for 2 min, followed by a final step at 72°C for 3 min) using the eubacterial primers 27F (AGA GTT TGA TCC TGG CTC AG) and 1492R (ACG GCT ACC TTG TTA CGA CTT) purchased from Integrated DNA Technologies, Inc. (USA). Cleanup of PCR products and DNA sequencing were performed at University of California Berkeley DNA Sequencing Facility. The PCR products were sequenced using the internal primers 27F and 1492R. Sequences were obtained by Sanger sequencing with ABI 3730XL DNA Analyzers (ThermoFisher, United States). Consensus sequences (1200–1400 base pairs) from forward and reverse sequences were generated using Geneious (version 9.1.3) and deposited in Genbank under the access codes listed in Table S2. The SILVA database was used for bacterial isolate classification. Sequence alignments, finding nearest neighbors, and phylogenetic tree reconstructions were performed in SILVA using SINA (v1.2.11) and RAxML. Phylogenetic tree visualization of the selected OTUs was performed using iTOL (https://itol.embl.de).

### Data analysis and statistics

Shannon’s diversity index (*H’*) and multivariate statistics were performed using the R package *vegan*. OTU distributions were transformed into relative abundances using the function *decostand*. These were subjected to Hellinger transformation before calculation of Bray-Curtis dissimilarity matrices comparing community composition between samples. Nonmetric multidimensional scaling (NMDS) using function *metaMDS* was performed using these dissimilarity matrices. A multivariate analysis of variance (MANOVA) model was implemented in the *vegan* function *adonis*. Analysis of similarity (ANOSIM) was carried out based on Bray-Curtis dissimilarities to evaluate the effect of C source and incubation time on community structure. We compared the relative abundance of taxa among the samples and selected ‘promoted’ OTUs using R software. Samples were compared by one-way ANOVA followed by the Dunnett’s test (*p* < 0.01) for multiple comparisons.

## Acknowledgement

This research by ENIGMA-Ecosystems and Networks Integrated with Genes and Molecular Assemblies (http://enigma.lbl.gov), a Scientific Focus Area Program at Lawrence Berkeley National Laboratory, is based upon work supported by the U.S. Department of Energy, Office of Science, Office of Biological & Environmental Research under contract number DE-AC02-05CH11231. The FRC groundwater sample was kindly provided by Terry C Hazen and Dominique C Joyner, Oak Ridge National Laboratory. We would also like to thank the MIT BioMicro Center for sequencing support. The sequencing efforts were funded by the National Institute of Environmental Health Sciences of the NIH under award P30-ES002109.

---

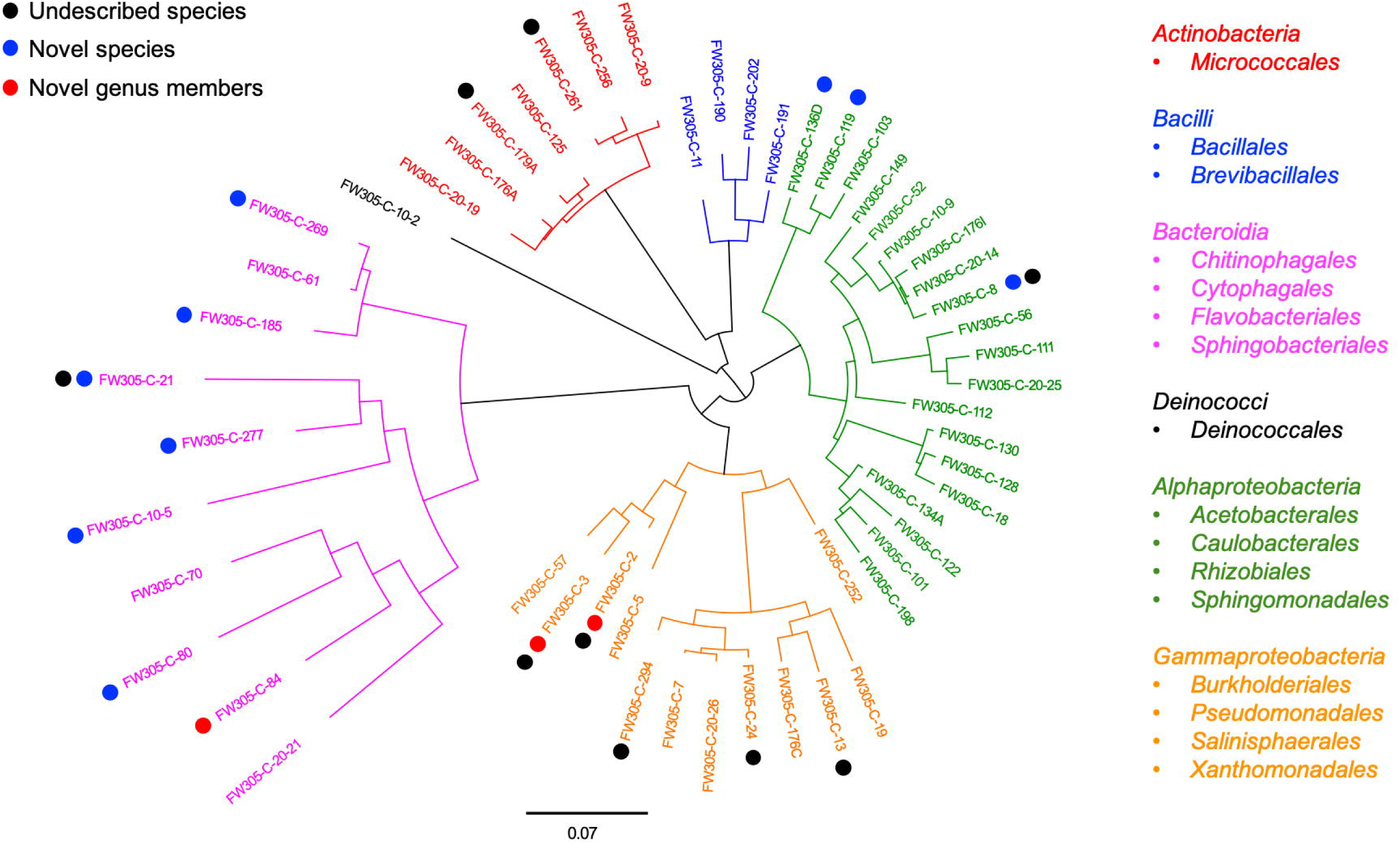

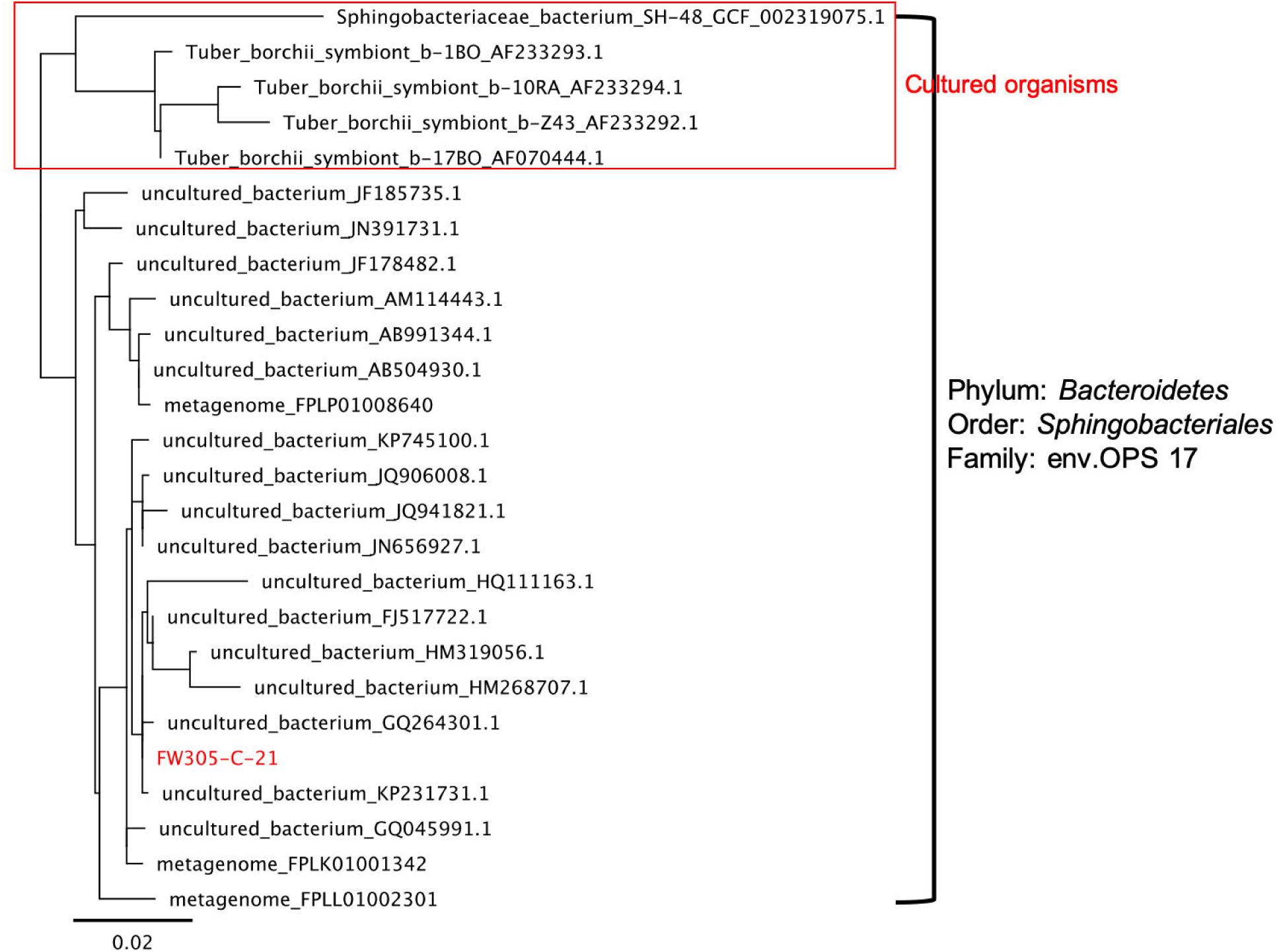

