## Supplementary figures and images for "Culturing of ‘Unculturable’ Subsurface Microbes: Natural Organic Carbon Source Fuels the Growth of Diverse and Distinct Bacteria from Groundwater"

### Figure S1

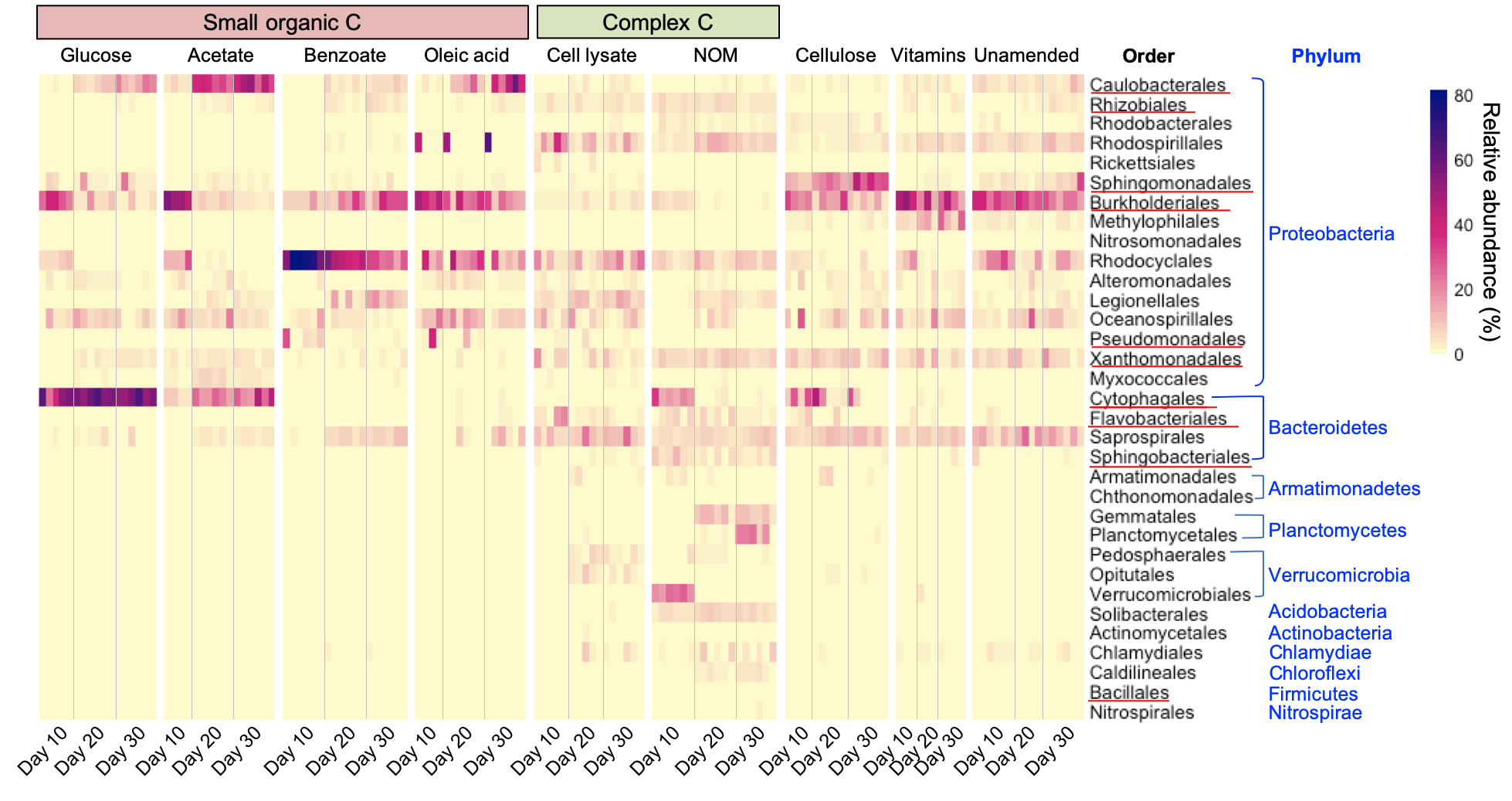
