## Supplementary Information for "Culturing of ‘Unculturable’ Subsurface Microbes: Natural Organic Carbon Source Fuels the Growth of Diverse and Distinct Bacteria from Groundwater"

**Table**

Table S1. Taxonomy of selected (one-way ANOVA with Dunnett’s multiple comparison test, *p*-value < 0.01) OTUs that were significantly promoted to grow in different C-amended enrichment cultures compared to unamended control, corresponding with OTUs illustrated in Figure 3A. Taxonomy was assigned using the SILVA database. Absence of an identified taxonomic level indicates no match with confidence > 0.5 returned from the SILVA database. Representative isolates (99-100% identity) obtained in this study are also included.

Table S2. The list of all 222 purified bacterial isolates obtained in this study.

Table S3. The list of representative isolates representing 54 distinct bacterial species, corresponding with isolates illustrated in Figure 4.

Table S4. Ingredients of vitamin stock solution.

| **Chemical** | **Concentration (mg/L)** |
| --- | --- |
| Vitamin B_1_ (Thiamine HCl) | 500 |
| Vitamin B_2_ (Riboflavin) | 500 |
| Vitamin B_3_ (Nicotinic acid) | 500 |
| Vitamin B_5_ (D-Pantothenic acid hemicalcium) | 500 |
| Vitamin B_6_ (Pyridoxine HCl) | 1000 |
| Vitamin B_7_ (Biotin) | 200 |
| Vitamin B_9_ (Folic acid) | 200 |
| Vitamin B_10_ (p-Amino benzoic acid) | 500 |
| Vitamin B_12_ (Cobalamin) | 10 |
| D,L-6,8-thioctic acid | 500 |
| **Total** | **4410** |

Table S5. Concentration of C substrate in stock solutions and microcosms.

| **C substrate** | **Conc. in stock solution** | **Final conc. of amended organic C in the microcosm (mg/L)** | **Note** |
| --- | --- | --- | --- |
| Glucose | 200 mM (36.0 g/L) | 144.1 | Nominal |
| Sodium acetate | 200 mM (16.4 g/L) | 48.0 | Nominal |
| Sodium benzoate | 50 mM (7.2 g/L) | 42.0 | Nominal |
| Oleic acid | 50 g/L (insoluble) | 382.7 (insoluble) | Nominal |
| Cellulose | 20 g/L (insoluble) | 88.9 (insoluble) | Nominal |
| Mixed vitamins | 4.4 g/L | 24.7 | Nominal |
| Bacterial cell lysate | TOC 2.67 g/L | 26.7 | Measured |
| Sediment NOM |  | 48.4 | Measured; NOM+groundwater background TOC |

Table S6. Primers used for 16S rRNA gene amplicon sequencing.

| **Primer name** | **Primer sequence (5’ -> 3’)** |
| --- | --- |
| PE16S_V4_U515_F* | ACACGACGCTCTTCCGATCTYRYRGTGCCAGCMGCCGCGGTAA |
| PE16S_V4_E786R* | CGGCATTCCTGCTGAACCGCTCTTCCGATCTGGACTACHVGGGTWTCTAAT |
| PE-III-PCR-F-### | AATGATACGGCGACCACCGAGATCTACACNNNNNNNNACACTCTTTCCCTACACGACGCTCTTCCGATCT |
| PE-IV-PCR-R-### | CAAGCAGAAGACGGCATACGAGATNNNNNNNNCGGTCTCGGCATTCCTGCTGAACCGCTCTTCCGATCT |

*Universal primer segments U515 and E786R were drawn from (Lane, 1991).

Lane, D.J. (1991) 16S/23S rRNA Sequencing. In: Stackebrandt, E. and Goodfellow, M., Eds., Nucleic Acid Techniques in Bacterial Systematic, John Wiley and Sons, New York, 115-175.

**Figure**

Figure S1. Relative abundance of enriched taxonomic order (>1% in any sample). Orders having representative isolates from this study are marked with red underlines.

Figure S2. Phylogenetic trees of selected isolates from candidate novel genera/species and undescribed species, and the most similar bacteria based on 16S rRNA genes. Scale bar indicates a change of a certain number per nucleotide. The 16S rRNA gene sequences were aligned using SINA against the SILVA alignment and the maximum likelihood tree was calculated using RAxML.

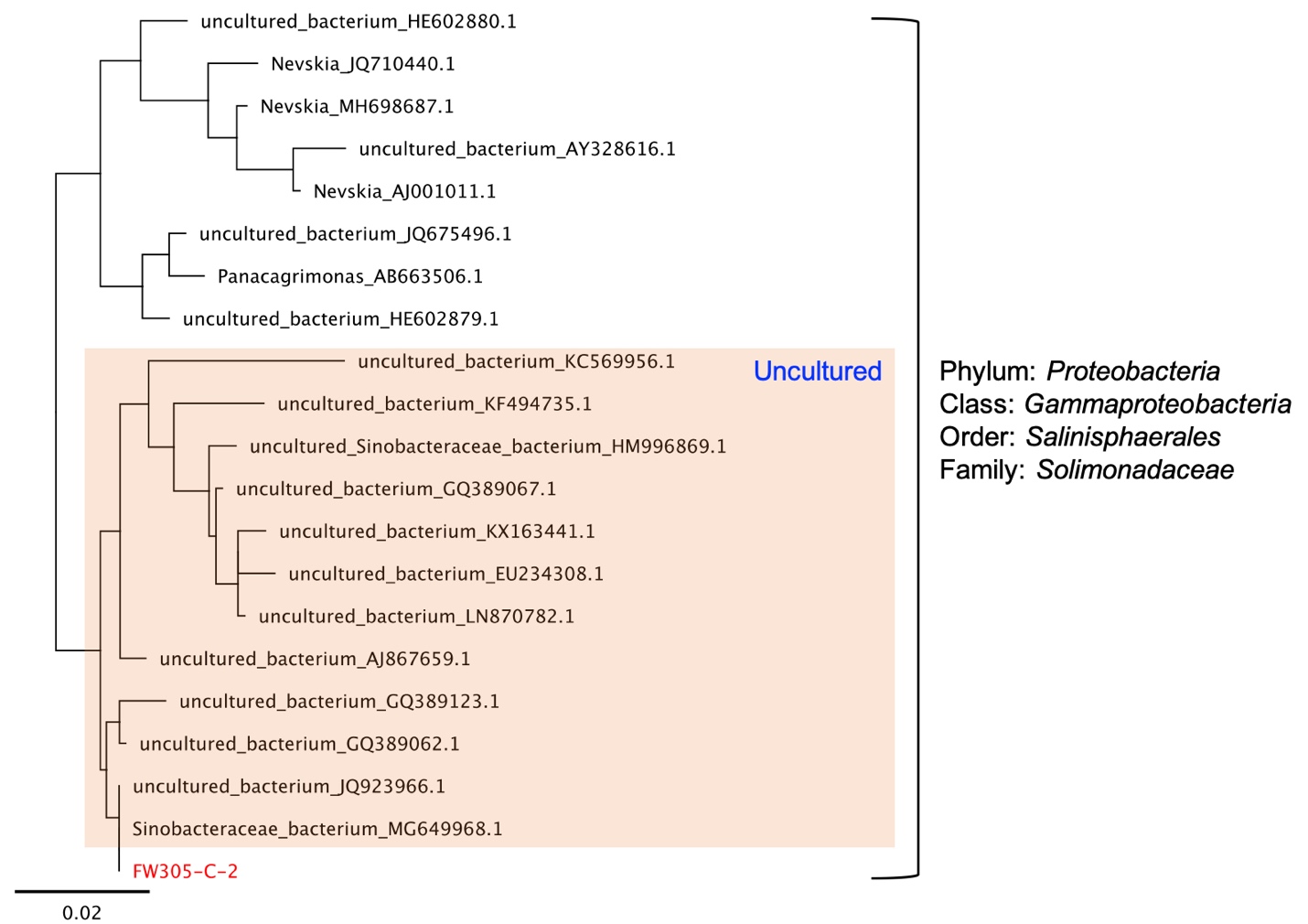

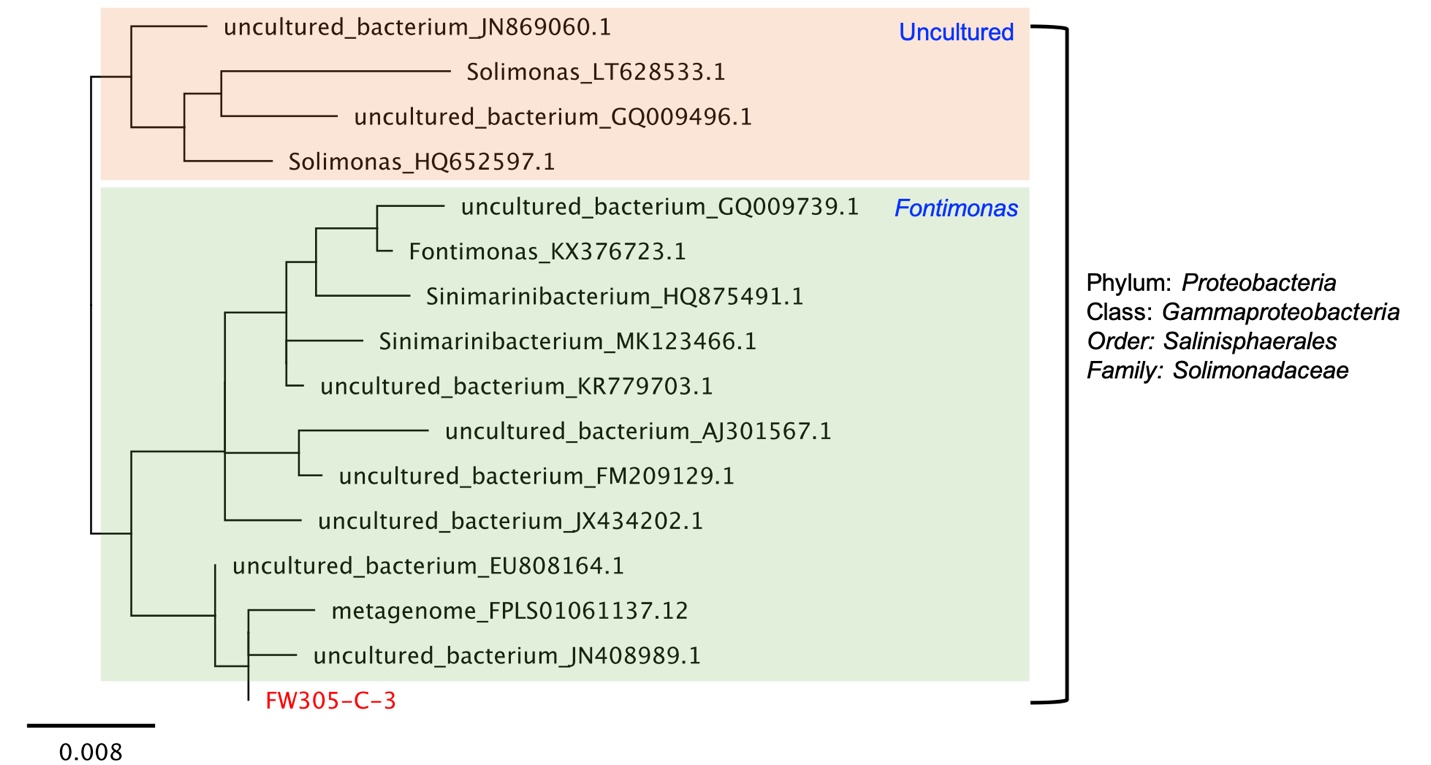

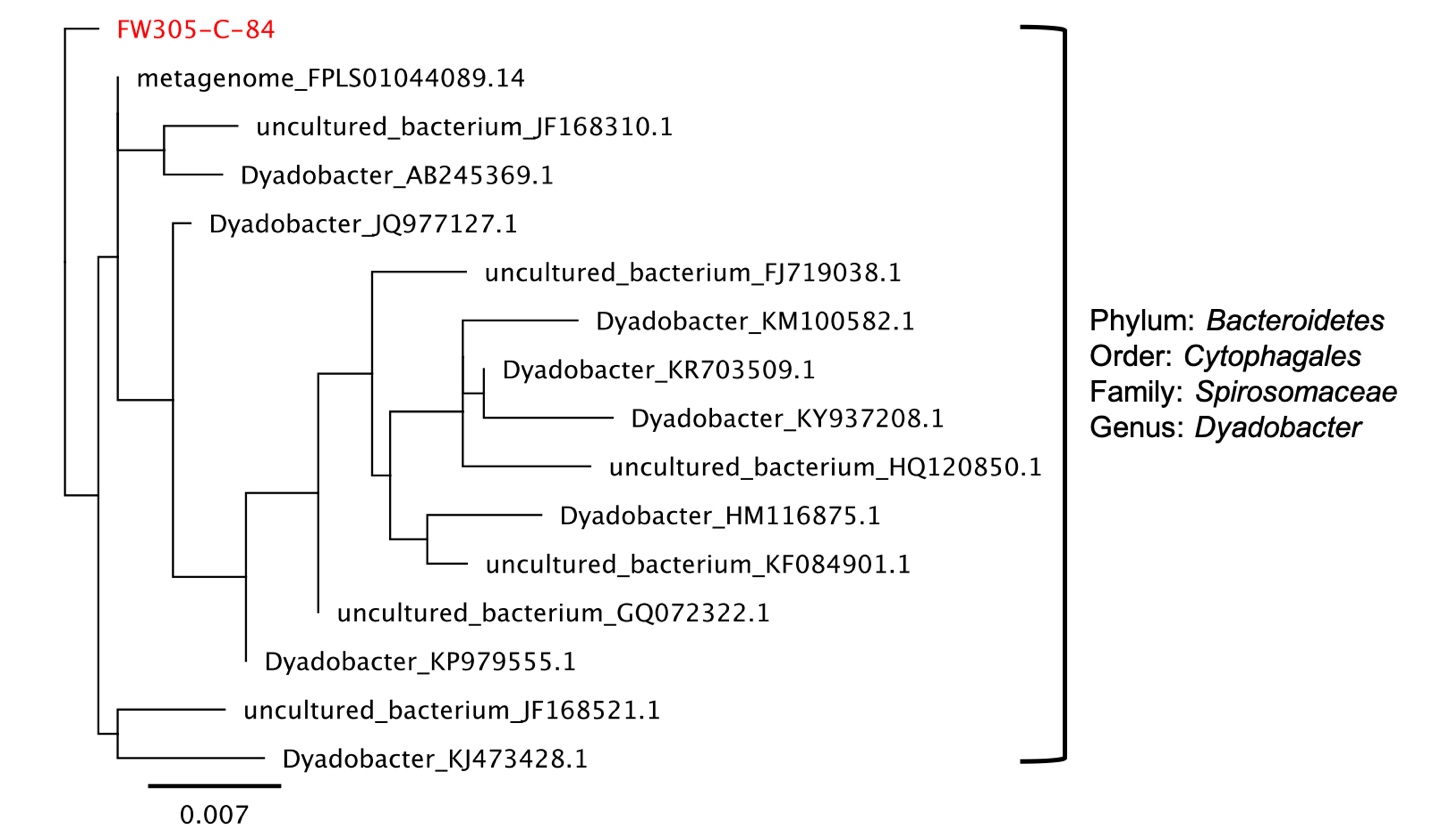

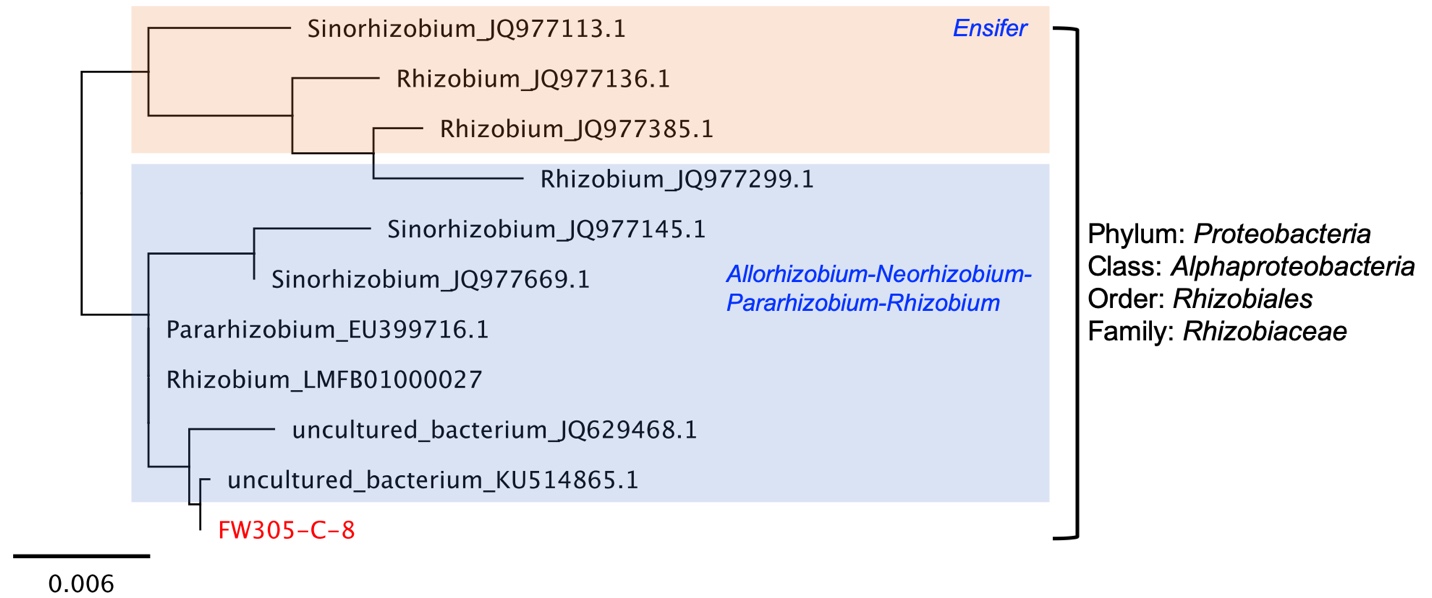

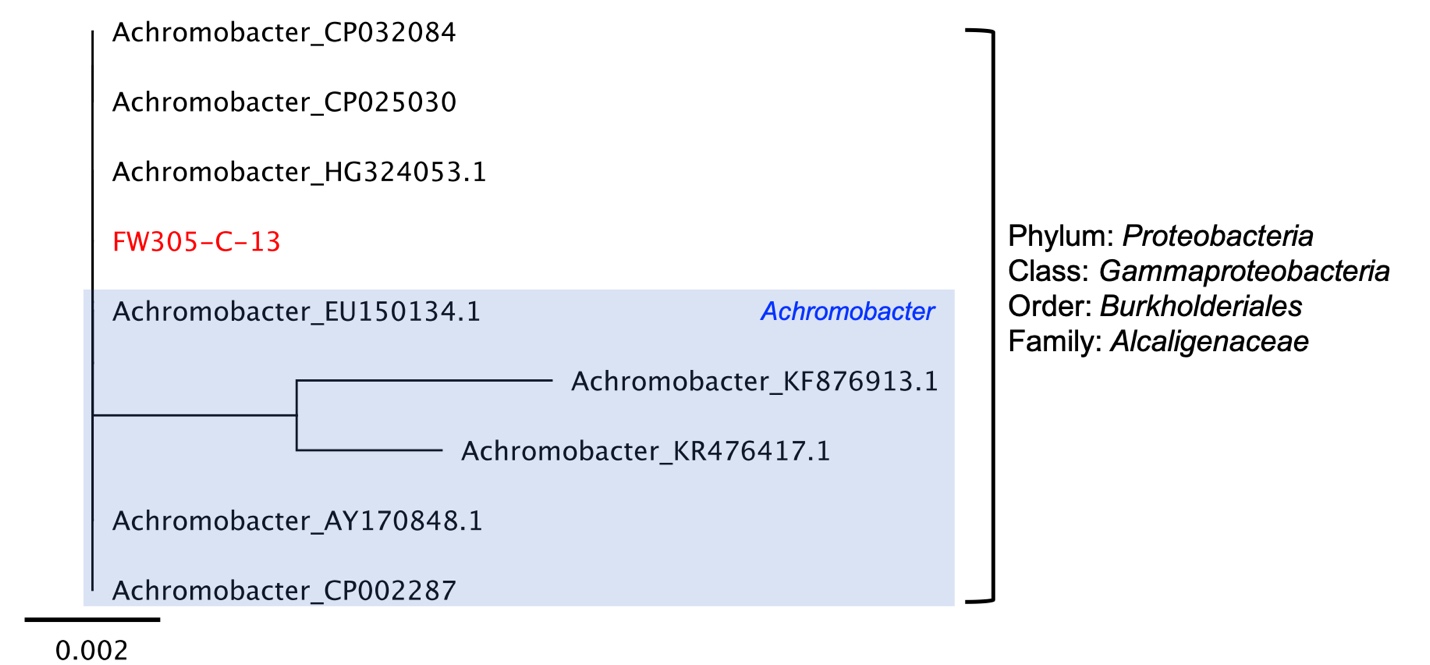

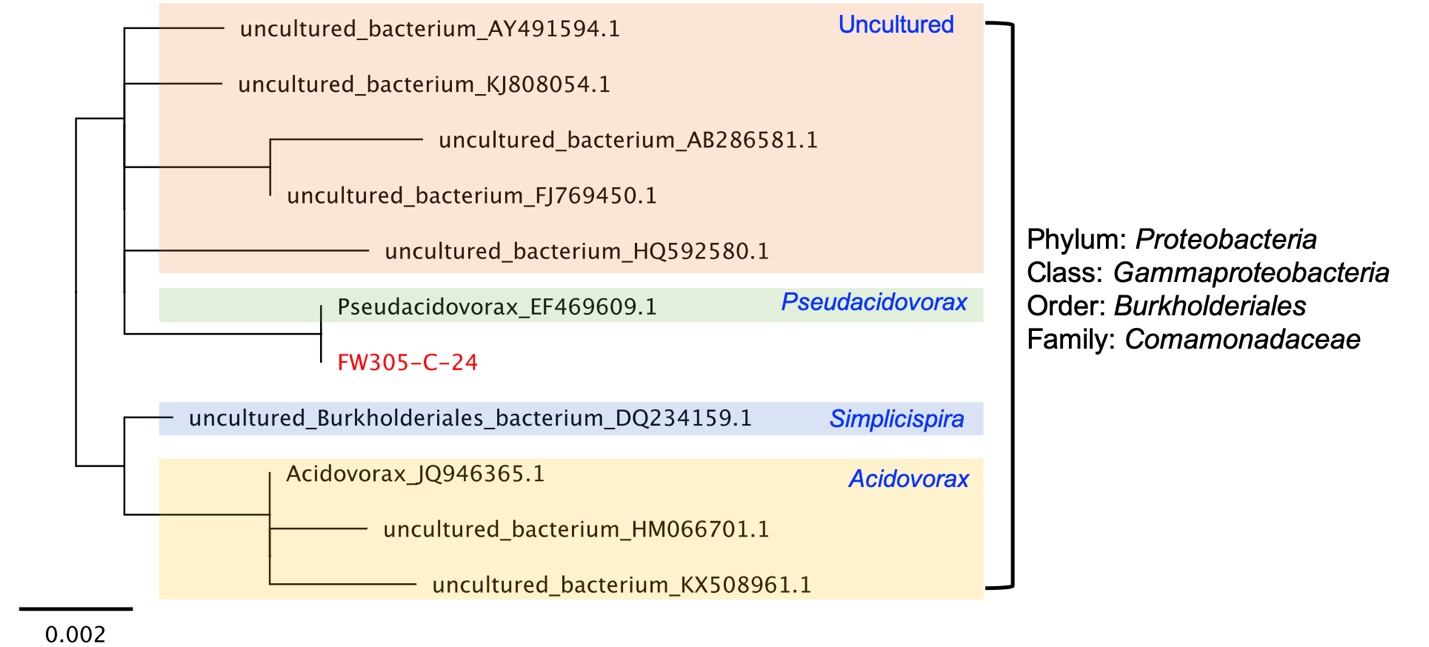

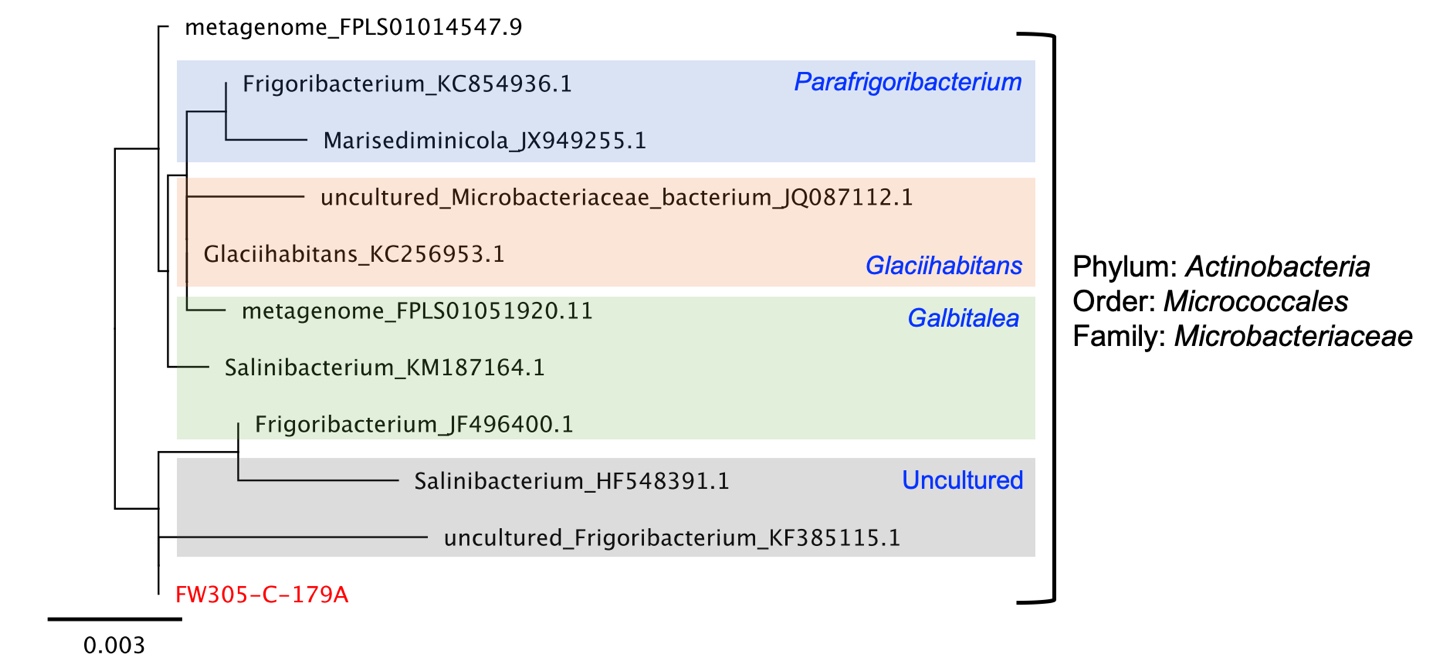

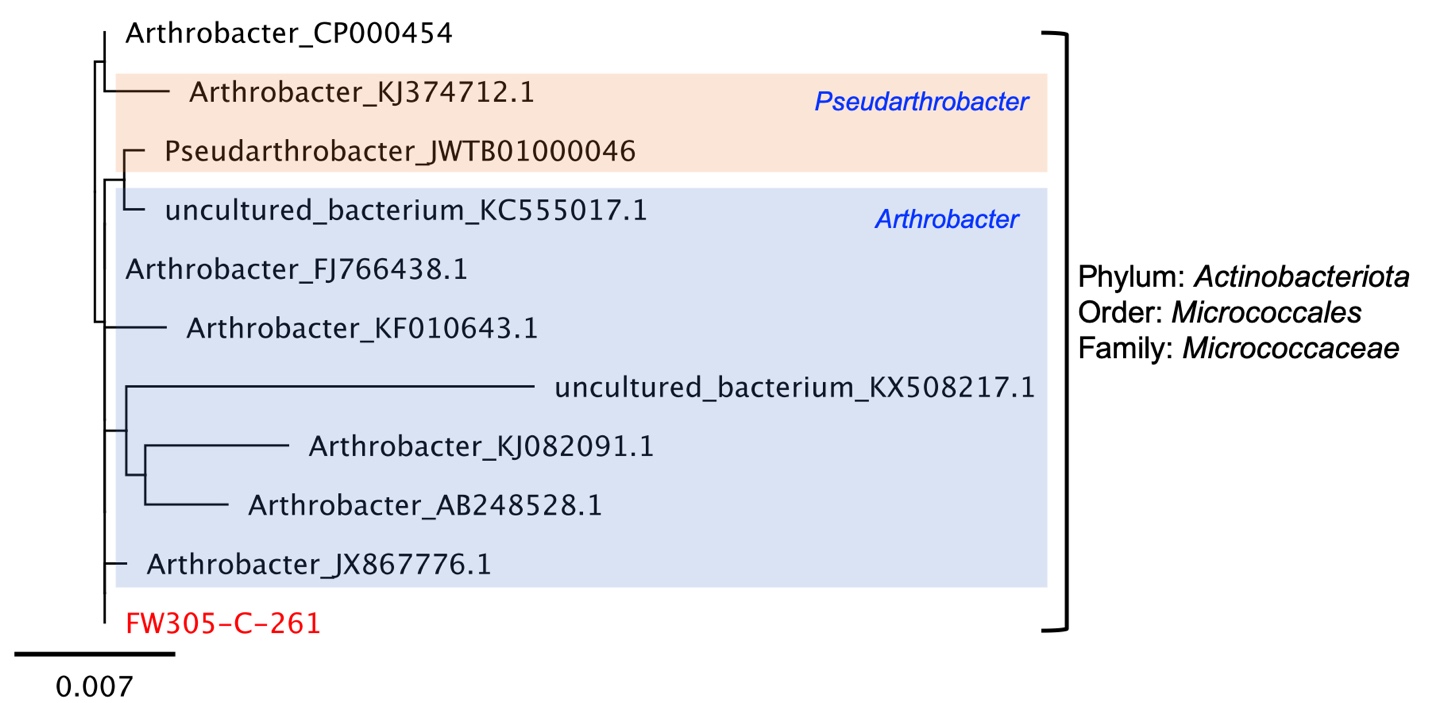

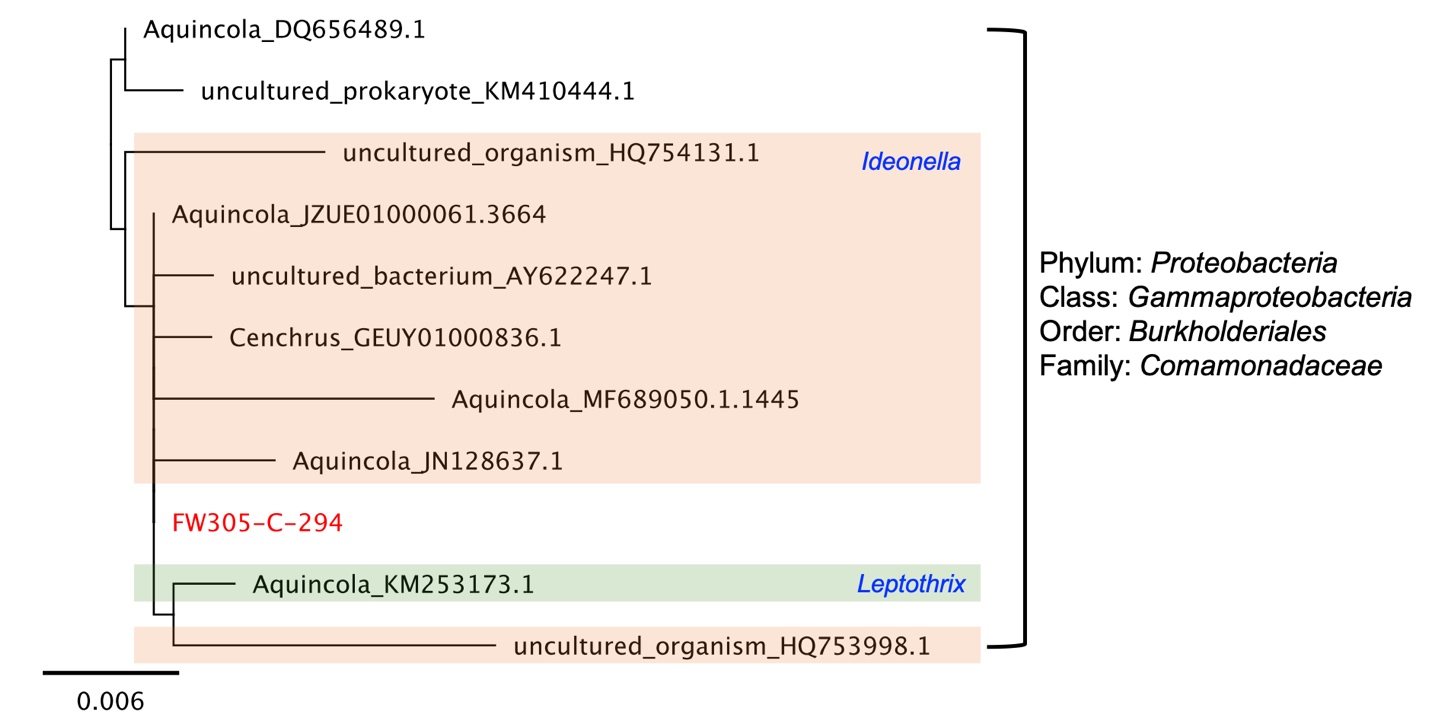
